# Nanoscale decompaction of nucleosomal DNA revealed through multi-color super-resolution microscopy

**DOI:** 10.1101/470823

**Authors:** Jason Otterstrom, Alvaro Castells Garcia, Chiara Vicario, Maria Pia Cosma, Melike Lakadamyali

**Author notes:** Co-first authors (Jason Otterstrom, Alvaro Castells Garcia). Co-last authors and corresponding authors: ML, MPC.

## Abstract

Chromatin organization plays an important role in regulating gene expression. Previously, we showed that chromatin is organized in the form of nucleosome groups or clutches. The size and nucleosome packing density of clutches decreased in hyperacetylated cells having more open chromatin. While hyperacetylation is thought to disrupt histone-DNA and inter-nucleosome interactions, its impact on higher order DNA compaction by groups of nucleosomes *in vivo* is not clear. To elucidate this question, we carried out two-color super-resolution imaging of histones and DNA in cells treated with the Histone Deacetylase (HDAC) inhibitor Trichostatin A (TSA). We showed that a lower percentage of DNA was associated to clutches in hyperacetylated cells, suggesting a decrease in nucleosome occupancy. We further identified the presence of “clutch” DNA within a nanoscale distance around the clutches. Upon histone hyperacetylation, the radius of the clutch DNA decreased leading to DNA release from the clutches, consistent with disruption of DNA-histone interactions. Finally, the most dramatic decompaction was observed for groups of clutches in close spatial proximity, suggesting that neighboring clutches influence each other’s DNA compaction.

**Summary:** Super-resolution imaging of histones and DNA reveals that DNA is compacted by groups of nucleosomes – clutches – at the nanoscale level and clutch compaction of DNA is affected by histone tail acetylation especially in highly folded regions containing several nearby clutches.

## Introduction

Each chromosome of interphase, eukaryotic nuclei occupies a specific nuclear area organized in chromosomal territories (CT) (Cremer and Cremer 2010; Andronov et al. 2016; Cremer et al. 2018). At the smallest of chromatin length scales, 146 base pairs of DNA wrap around an octamer of histone proteins, forming the nucleosome. Nucleosomes arrange as beads on a string along the DNA forming a fiber that is roughly 10 nm in diameter. Through the linker histone H1, the 10nm fiber further compacts into higher order structures (Thoma et al. 1979). *In vitro* reconstitution studies showed that nucleosomes form a very ordered zig-zag or solenoid-like organization compacting chromatin into a 30 nm fiber. However, the existence of this 30 nm fiber *in vivo* has been long debated and how the chromatin compacts and folds into higher order structures remains unclear. Several recent studies suggested that the organization of nucleosomes and higher order chromatin folding is much more heterogeneous than the regular 30 nm fiber (Fussner et al. 2012; Joti et al. 2012; Maeshima et al. 2010; Quenet et al. 2012; Ricci et al. 2015; Eltsov et al. 2008; Nishino et al. 2012; Ou et al. 2017). The emerging picture suggests that chromatin is largely comprised of 10 nm fibers with different levels of compaction. In line with these recent studies, our previous work using super-resolution microscopy demonstrated that nucleosomes form heterogeneous groups, termed nucleosome clutches, with clutch density and area correlating with the level of chromatin compaction and inversely correlating with the cell pluripotency grade (Ricci et al. 2015).

Chromatin compaction and folding is further shaped by histone post-translational modifications. The core histones, and in particular H3 and H4, have N-terminal protein tails protruding from the nucleosomes, which can be covalently modified at different residues (Bannister and Kouzarides 2011; Rothbart and Strahl 2014). The tails can be methylated, acetylated, phosphorylated, or ubiquitinated, among other modifications. The combination of these different modifications has been defined as a “histone code” (Jenuwein and Allis 2001). The histone code of nucleosomes positioned at promoter regions biases DNA accessibility to transcription factors and transcription machinery, and is an important regulator of gene activation and repression (Li et al. 2007). Specifically, lysine acetylation neutralizes lysine positive charges. This neutralization results in a reduction of the electrostatic attraction between the histone tails and the negative charged DNA. It has thus been postulated (Eberharter and Becker 2002; Verdone et al. 2005) that histone acetylation results in a more “open” chromatin fiber with the DNA more loosely attached to the nucleosomes. In addition, all-atom molecular dynamics simulations suggested that acetylation can disrupt inter-nucleosome interactions leading to unfolding and de-compaction of the chromatin fiber (Collepardo-Guevara et al. 2015). This chromatin opening leads to an increased accessibility of the transcription factors and RNA Polymerase II holoenzyme and therefore to transcription activation (Eberharter and Becker 2002; Verdone et al. 2005). While previous work using *in vitro* reconstituted chromatin and all-atom molecular dynamics simulations lead to a molecular model of acetylation-induced chromatin fiber decompaction, how acetylation impacts chromatin structure *in vivo* is less clear.

To address the *in vivo* mechanisms of how groups of nucleosomes compact DNA and how this compaction is affected by hyperacetylation, we combined 3D STORM and PAINT super-resolution microscopy to image DNA simultaneously with histones at the nanoscale level *in situ* within human fibroblast cells. Quantitative spatial analysis of DNA and histone co-organization showed that a larger percentage of DNA is free of nucleosome clutches in hyperacetylated cells. We further identified what we refer to as a “clutch” DNA present within a small nanoscale area around the nucleosome clutches, whose compaction is influenced by the nucleosome clutch. This clutch DNA partially decompacted after histone acetylation. Finally, we observed that the clutch DNA decompaction is largely influenced by the presence of nearby clutches. Overall, our results quantify the structural reorganization and DNA decompaction induced by histone tail acetylation at the nanoscale level *in vivo*, complementing the molecular understanding previously inferred through *in vitro* experiments and mesoscale modeling.

## Results

To visualize DNA organization using Stochastic Optical Reconstruction super-resolution microscopy (STORM), human Fibroblast cells (hFb) were labeled with a nucleotide analog 5-Ethynyl-2′-deoxycytidine (EdC) (see Materials and Methods)(Raulf et al. 2014; Zessin et al. 2012). EdC was provided at high concentration (5 μM) and for long enough time period (4 days, see Materials and Methods) to ensure dense DNA labeling and enable visualization of global DNA organization. 67 ± 10% of cells were positive for EdC after this treatment **(Figure S1 A)** and EdC labeling had minimal impact on cell cycle at the concentration and incubation time used (**Figure S1 B**). Subsequent fixation and click chemistry with AlexaFluor647 allowed 3D super-resolution imaging of DNA structure (**Figure 1A, upper**). Super-resolution images of DNA were segmented using Voronoi tessellation (Andronov et al. 2016; Levet et al. 2015) and color coded according to the inverse of the area of Voronoi polygons (**Figure 1B upper and 1C, left**). The Voronoi polygon area is inversely related to DNA density, with higher density regions corresponding to smaller Voronoi polygons and vice versa. Although localizations were extracted in 3D, the analysis was performed on a 2D projection of a thin, 120 nm slice through the center of the nucleus, which contained approximately 33% of all localizations (**Figure S2**). The 120 nm z-thickness was selected to include a sufficiently high density of localizations for structural analysis while minimizing 2D projection effects. This volume is also in a similar range as recent electron tomographic reconstruction of DNA organization (Eltsov et al. 2008; Ou et al. 2017). Compact, heterochromatic regions typically located at the nuclear edge and surrounding the nucleoli in human fibroblasts were characterized by high DNA density in the super-resolution images (**Figure 1B, upper**), suggesting that DNA density can, indeed, be a proxy of chromatin compaction and heterochromatin. Treatment with the Histone Deacetylase (HDAC) inhibitor Tricostatin A (TSA) has been shown to lead to hyperacetylation of histone tails, chromatin decompaction and increased transcriptional activity (Toth et al. 2004). Consistent with these previous results, TSA treatment induced a decrease in the median DNA density using the Voronoi tessellation (**Figure 1D**). Therefore, DNA density in the Voronoi segmented super-resolution images is, indeed, indicative of the level of DNA compaction.

**Figure 1.**
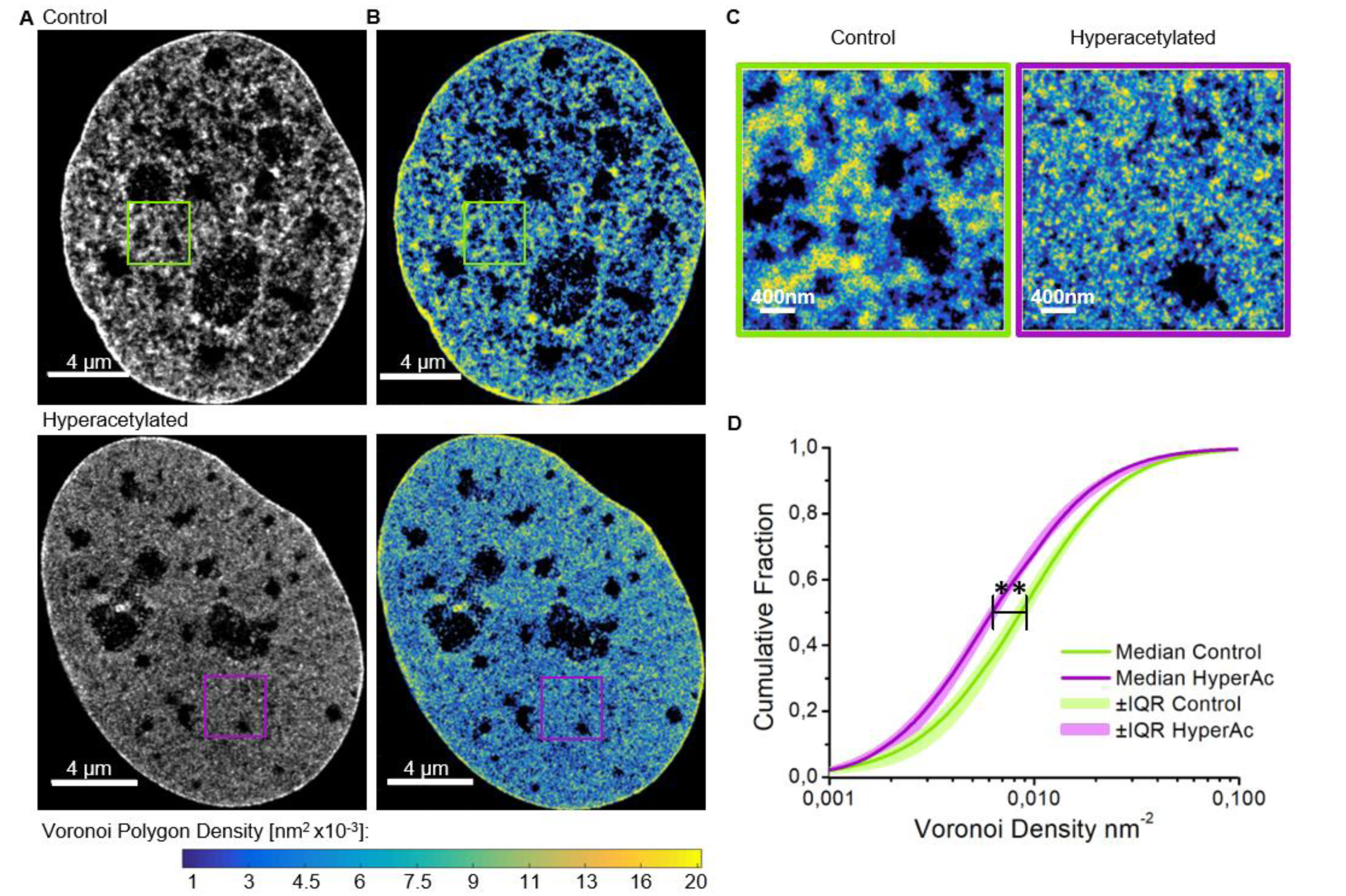
EdC labeling enables super-resolution imaging of DNA structure. **(A)** Cropped nuclear super-resolution images of EdC labeled DNA in control (upper) and TSA-treated (lower) human BJ fibroblast cells. **(B)** Super-resolution Voronoi tessellation image of DNA in control (upper) and TSA-treated (lower) fibroblasts. Voronoi polygons are color-coded according to the density (inverse of the polygon area) following the color scale bar (from 0.02 nm2 in yellow to 0.001 nm2 in blue; the largest 0.5% of the polygons are colored black). **(C)** A zoom up of the region within the squares in (B). **(D)** Cumulative distribution of the Voronoi Polygon densities in control (green) (N=6 cells) and TSA-treated (magenta) (N=9 cells) fibroblasts. The light colors show the interquartile range (25-75 percentiles) and the thick, dark lines show the median values; stars indicate statistical significance of the separation between the median of the medians according to Kolmogorov Smirnoff test with p = 0.0022.

Previously, we showed that nucleosomes are organized in heterogeneous groups, which we termed nucleosome clutches, and demonstrated that nucleosome clutch size and density decreased in TSA-treated human fibroblasts (Ricci et al. 2015). To gain further insight into chromatin decompaction mediated by hyperacetylation, we aimed to carry out two-color super-resolution imaging of DNA and histone H2B to simultaneously visualize remodeling of nucleosome clutches together with their associated DNA. Since the click chemistry used to label DNA offers a limited choice for reliable super-resolution compatible, photoswitchable fluorophores (mainly AlexaFluor647), we aimed to identify a second fluorophore with a different emission wavelength to carry out the immunofluorescence labeling of histones. To this end, we first tested a wide range of fluorophores previously reported to be compatible with super-resolution microscopy (Dempsey et al. 2011) and compared the histone clutch structure to those obtained by the best-performing AlexaFluor647 (**Figure S3A**). The histone data were analyzed using our previously developed distance-based cluster identification algorithm and the localizations were segmented into nucleosome clutches ((Ricci et al. 2015), **Figure S3B**).

Detailed analysis of these data showed that most fluorophores gave rise to sparse appearance of the clutches in the rendered images compared to those obtained by AlexaFluor647 (**Figure S3A and C**). Indeed, the nuclear area occupied by fluorophore localizations was significantly lower for Cy3B, AlexaFluor568, Atto488 and AlexaFluor750 compared to AlexaFluor647 (**Figure S3C**). Nucleosome clutches in STORM images of AlexaFluor647 typically cluster together, with multiple clutches in close spatial proximity of one another (**Figure S3B**), likely corresponding to the higher order folding of the chromatin fiber. This result is consistent with recent super-resolution imaging of single chromosomes, showing multiple DNA nanoclusters in close proximity (Fang et al. 2018). However, nucleosome clutches imaged with alternative fluorophores were mostly isolated in space and had significantly fewer neighboring clusters (**Figure S3D, E**). We attribute these effects to the poor photoswitching properties of the alternative dyes, in particular in the nuclear environment. Indeed, the localization density per frame was substantially higher for AlexaFluor647, giving rise to a much higher cumulative number of localizations in the same image acquisition time compared to other fluorophores (**Figure S3F**). Accordingly, the final image resolution computed using the Fourier Ring Correlation (FRC) analysis (Banterle et al. 2013) (**Figure S3G**) was significantly improved for super-resolution images of histones acquired using AlexaFluor647 compared to other fluorophores (**Figure S3H**).

To circumvent this problem, we next combined two different super-resolution imaging modalities: STORM to image DNA and PAINT to image histones. The DNA-PAINT (Schnitzbauer et al. 2017), referred here simply as PAINT for clarity, does not rely on the use of imaging buffers and fluorophore photoswitching. Instead, the on-off blinking of fluorophores is achieved by the transient, reversible binding of fluorophore labeled “imager oligo strands” to the complimentary “docking oligo strands”. The docking strands are conjugated to a secondary antibody (Schnitzbauer et al. 2017), which is used for immunostaining and hence imaging a target protein. Hence, a wide range of fluorophores can be used for PAINT imaging. Our results showed that PAINT gave rendered super-resolution images of H2B that were both qualitatively and quantitatively comparable to those obtained by STORM imaging using AlexaFluor647 (**Figure S3A, C-H**). Importantly, the number of localizations per nucleosome clutch decreased after TSA treatment when H2B was imaged with PAINT or with STORM using AlexaFluor647 (**Figure S3I**). This result is consistent with nucleosomal decompaction upon hyperacetylation and in accord with our previous results (Ricci et al. 2015). To combine the two super-resolution modalities for multi-color imaging, we designed an analysis workflow that accounts for the differences in acquisition time needed for STORM and PAINT (see Materials and Methods). Finally, to correct for drift and to align the two images in 3D, we used fiduciary markers that were internalized into cells prior to fixation (**Figure S4A**). To measure the residual registration error, we imaged H2B via PAINT in two colors (560nm and 647nm laser excitation) to obtain mock two-color images of the same structure **(Figure S4B)**. Nearest neighbor distances between clutches present in each color allowed calculation of a 3D registration error of 31 ± 15 nm (**Figure S4C**).

Two-color super-resolution images of histones and DNA in control cells revealed, as expected, a similar pattern of heterogeneous labeling, showing regions of the nucleus enriched in and depleted of chromatin (**Figure 2A, Figure S5**). To further quantify the spatial relationship between histones and DNA, we first segmented the nucleosome clutches and determined their center position. The center position of each clutch was then used as seeds for Voronoi tessellation. This analysis divided the nuclear space into polygons, each polygon surrounding one clutch center (**Figure 2B**). The DNA localizations falling within the Voronoi polygon area of a particular clutch center were then considered for further analysis. We next determined the DNA localization density within circles having a range of search radii from the clutch center. The circles were bounded by the edges of each clutch’s Voronoi polygon (i.e. if the search radius was larger than the polygon, then the edges of the polygon were used as the boundary) (**Figure 2B**). This approach guaranteed that each DNA localization is assigned to only one nucleosome clutch. The DNA density around nucleosome clutches in TSA treated cells was lower compared to control cells over the full range of search radii utilized, indicating decompaction of clutch associated DNA after hyperacetylation (**Figure 2C**).

**Figure 2:**
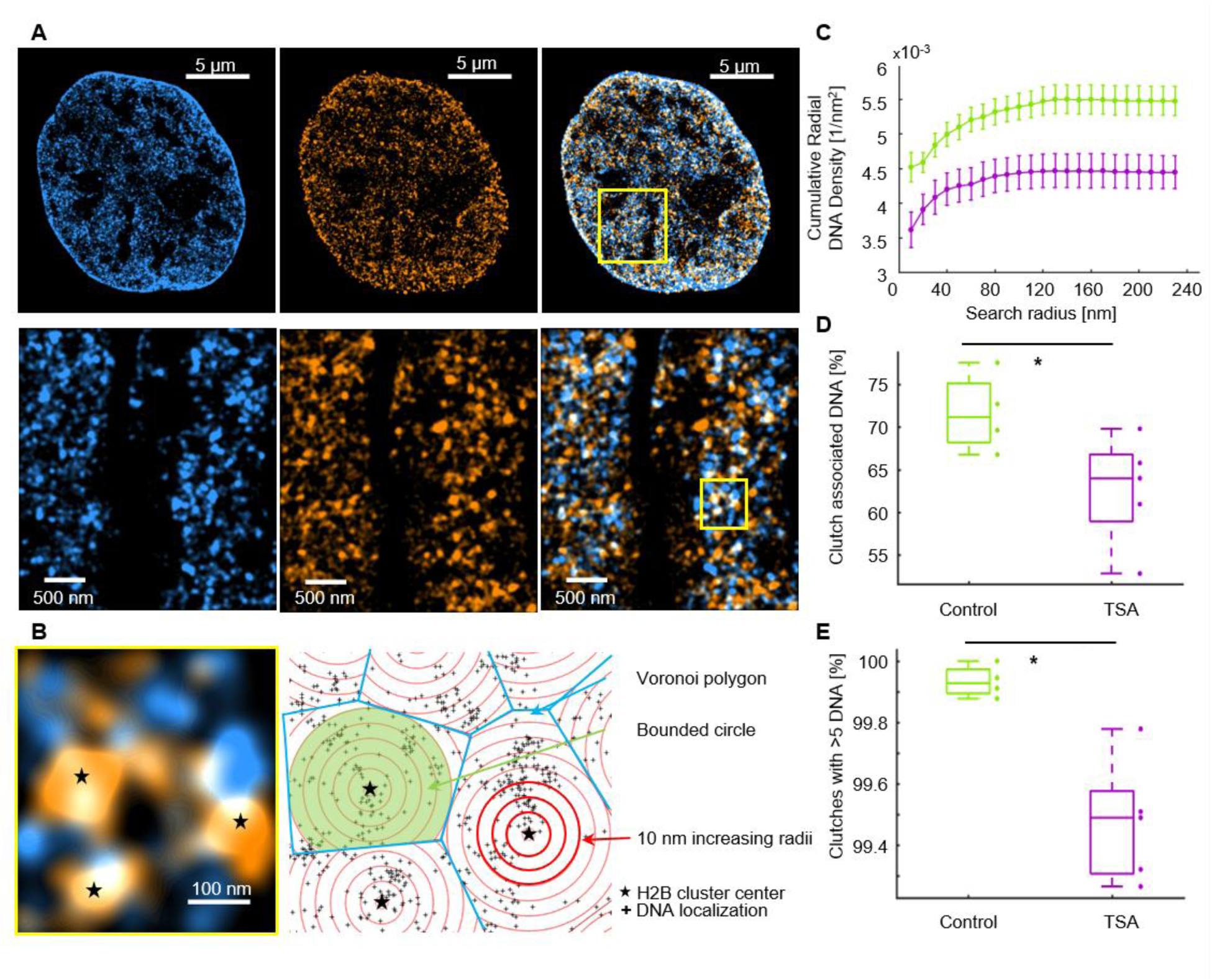
DNA co-localizes with H2B to a lesser extent in TSA-treated cells compared to untreated cells. **(A)** Cropped nuclear super-resolution image of EdC-labeled DNA (cyan) and PAINT image of H2B labeled with anti-H2B antibodies (orange) and the overlay. A zoom of the region inside the yellow box is shown. **(B) (left)** A zoom of the region shown inside the yellow square, **(right)** scheme of the analysis of clutch-bound DNA. The centers of H2B clusters (stars) are the seeds for the Voronoi polygons (blue) inside which the DNA localizations (black dots) are distributed. Overlaid on top are concentric circles whose radii increase by 10 nm steps. **(C)** Cumulative DNA density inside circles of increasing search radii in untreated (green) and TSA-treated (magenta) cells. The dots correspond to the mean, the bars correspond to the standard deviations: the lines connect sequential points and are guides to the eye. **(D)** Percentage of DNA localizations associated to clutches in wild-type (N=4, green) and TSA-treated (N=5, magenta) fibroblasts for a circle of radius 120 nm bounded by Voronoi polygons (p-value 0.0441). **(E)** Percentage of H2B clutches associated to DNA in wild type (green) and TSA-treated (magenta) cells (p-value 0.0168). Stars indicate statistical significance according to an unpaired two-tailed t-test with Welch correction.

We next used a bounded circle with a search radius of 120 nm and categorized the DNA localizations falling within the circle as ‘clutch-associated’ and those falling outside as ‘clutch-free’ DNA. The 120 nm radius was selected based on the fact that the median DNA density reached a plateau and was stable at this radius for both untreated and TSA treated cells (**Figure 2C**). This radius also corresponded to about 2.5 times the average standard deviation of the localizations belonging to the H2B clutches, therefore encompassing ~98% of the nucleosome clutch area. This co-localization analysis showed that a higher proportion of DNA was nucleosome clutch-associated in wild type Fibroblast cells compared to TSA treated Fibroblast cells (71.7±2.3% in WT Fb and 62.7±2.9% in TSA treated Fb, n=4 and n=5 cells, respectively, **Figure 2D**). These observations are consistent with the results of our previous polymer based modeling of DNA occupancy from single color images of nucleosome clutches, which also showed a decrease in percentage of DNA occupied by nucleosomes after TSA treatment (Ricci et al. 2015). On the other hand, a large percentage (~99%) of histones co-localized with DNA in both conditions, suggesting that most histones are chromatin associated as expected (**Figure 2E**).

To get a more detailed understanding of how the nucleosome clutches compact DNA in untreated and TSA-treated cells, we next analyzed the DNA density within 10 nm thick rings of increasing radii (**Figure 2B**). The ring radius was iteratively increased by 10 nm intervals while keeping the thickness constant and hence moving the ring further away from the clutch center (**Figure 2B**). We calculated the DNA density within each 10 nm ring from the DNA localizations falling within the ring area. We then created a “similarity matrix” by comparing the DNA density among rings of different radii (**Figure 3A**). In untreated cells, there was low similarity in DNA densities of rings with radii ranging from 10-70 nm (**Figure 3A, left, cyan bars**), suggestive of variation in the DNA compaction level over this length scale. Search radii above 70 nm, on the other hand, showed high similarity in their DNA density. Therefore, within a 70 nm distance from the clutch center there is large heterogeneity in DNA compaction whereas at larger distances, DNA compaction is unchanged. These results suggest that nucleosome clutches compact DNA within a 70 nm radius from their center in untreated cells. We call the DNA that falls within the low similarity region of the matrix “clutch” DNA: *i.e*. DNA associated to a clutch, which includes both the DNA wrapped around individual nucleosomes as well as the linker DNA in between neighboring nucleosomes. Interestingly, in TSA-treated cells the clutch DNA radius decreased to ~40 nm (**Figure 3A, right, red bars**). This result suggests that hyperacetylation leads to a de-compaction of DNA that lies within a zone of 40-70 nm from the clutch center and it is consistent with the fact that the average clutch size in TSA treated cells is smaller (Ricci et al. 2015). Instead, in untreated fibroblasts the clutch DNA is not only denser but it also spans a larger distance from the clutch center. These results are in agreement with a disruption of histone-DNA interactions for nucleosomes having acetylated tails.

**Figure 3.**
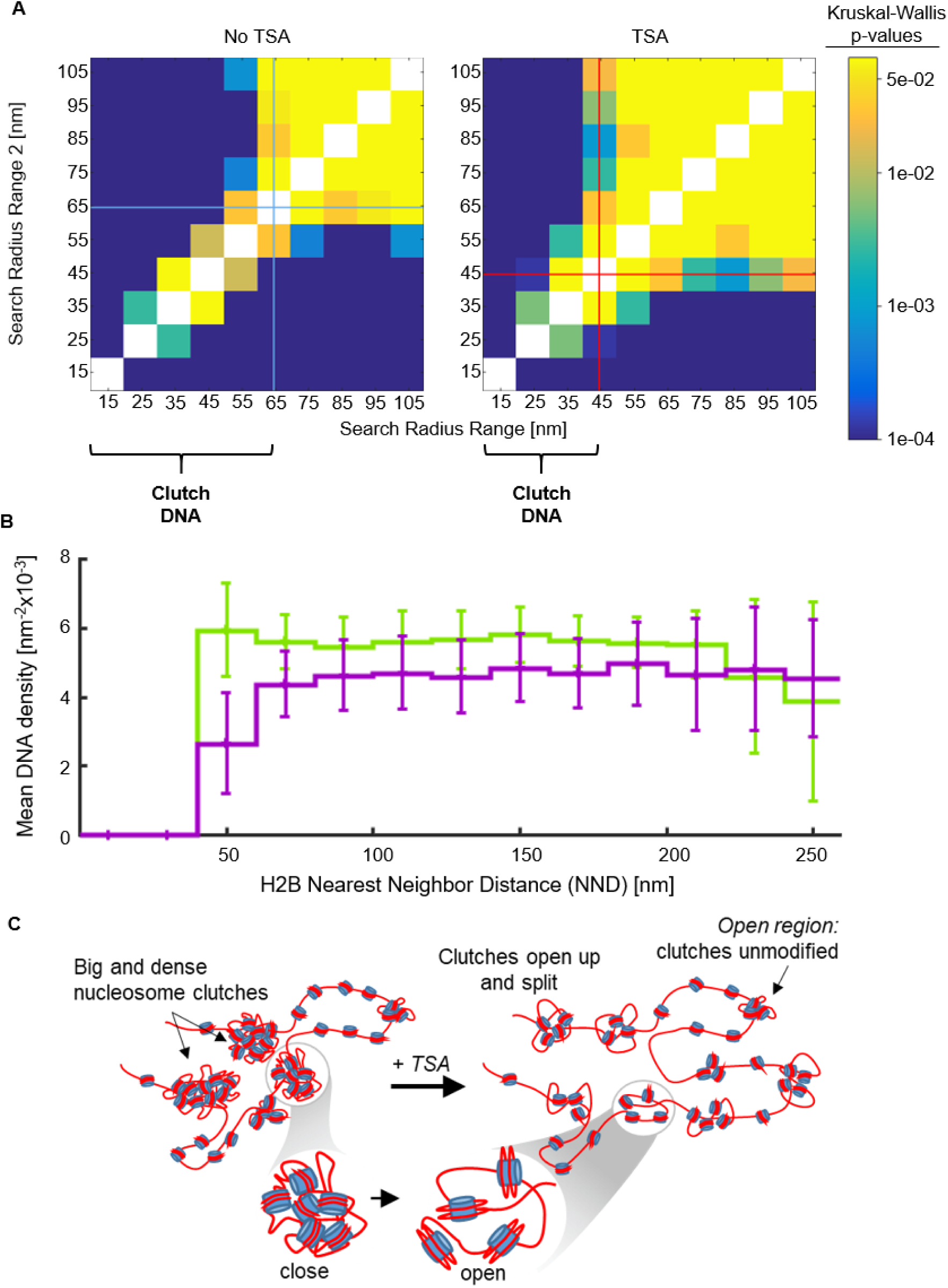
Nucleosome clutch associated DNA undergoes decompaction in TSA-treated cells: **(A)** Similarity matrix for untreated (left) and TSA-treated (right) cells showing the level of similarity in DNA density within 10 nm discs of increasing radii. The similarity was calculated as a p-value from Kruskalwallis test and is shown as a color coding corresponding to the color scale bar (from p=0.0001 in blue to p=0.05 in yellow). The diagonal was set to white and not calculated. Cyan and red lines show the boundary of a switch from low to high similarity in untreated and TSA treated cells, respectively **(B)** Mean DNA density ± standard deviation as a function of clutch nearest neighbor distance (NND) for untreated (green) and TSA-treated (magenta) cells calculated for a circle of radius 70 nm. The bars show the standard deviation. **(C)** Cartoon model of nucleosomal DNA decompaction upon hyperacetylation. Nucleosome clutches become smaller, likely due to a combination of large clutches opening and splitting and nucleosome loss. Clutches also compact “clutch” DNA to a lesser extend in hyperacetylated cells. These changes are particularly prominent in highly folded regions of the chromatin containing multiple nucleosome clutches in close proximity, suggesting that clutches may influence DNA compaction of neighboring clutches.

In human fibroblasts, clutches further clustered in close spatial proximity of one another, forming “islands” containing several clutches organized into higher order structures, which were hundreds of nanometers in size (**Figure S3B**). We reasoned that these regions might correspond to the higher order folding of the chromatin fiber, bringing multiple clutches into close proximity to further compact DNA. We next wondered how the DNA packing changes upon hyperacetylation in relation to these folded regions of the chromatin. We compared the DNA density falling within a radius of 70 nm from the clutch center (the clutch DNA for untreated cells) as a function of the nearest neighbor distances (NNDs) between clutches (**Figure 3B**). Surprisingly, the largest difference in DNA density between untreated and TSA treated cells was observed for clutches having an NND of ~50 nm or smaller (**Figure 3B**). These results suggest that the largest decompaction due to hyperacetylation happens in folded chromatin regions containing high-order structures, where decompacted DNA from one nucleosome clutch may influence the DNA packing of a nearby, neighboring clutch (**Figure 3C**), thus changing the DNA density of multiple clutches. This decompaction might be due to the presence of a high density of nucleosome tails that become hyperacetylated in these regions, thus generating a large amount of repulsive charges that more strongly disrupt DNA-histone interactions leading to higher DNA decompaction. On the other hand, the more unfolded regions corresponding to more isolated clutches do not undergo major modifications, likely since chromatin in these regions is already in an open state (**Figure 3C**).

## Discussion

Visualization of DNA fibers at nanoscale resolution within an intact nucleus can potentially reveal details about how chromatin organization relates to gene activation, information that biochemical or genetic approaches can only indirectly infer (Rowley and Corces 2018). In recent years, super-resolution microscopy has been revealing the spatial organization of chromatin and transcriptional machinery in mammalian and bacterial cells at length scales that were previously inaccessible to light microscopy and that are highly relevant to gene activity (Beliveau et al. 2015; Boettiger et al. 2016; Ricci et al. 2015; Wang et al. 2016; Cho et al. 2018; Cisse et al. 2013; Weng and Xiao 2014). Labeling of specific genomic regions using Oligopaint technology and imaging with super-resolution (OligoSTORM) uncovered the higher order packing of genomic regions having different epigenetic states (Beliveau et al. 2015; Boettiger et al. 2016; Wang et al. 2016). By using super-resolution microscopy, we previously revealed the organization of the chromatin fibers by visualizing nucleosome clutches (Ricci et al. 2015). More recently, pulse-chase EdU labeling and click chemistry followed by super-resolution microscopy was used to label the DNA of individual chromosomes (Fang et al. 2018). These images showed that DNA formed nanodomains, which were further clustered in close spatial proximity, an organization similar to nucleosome clutches that also cluster in close proximity. In addition, multi-color imaging of DNA and histone modifications revealed that active histone marks formed small, disperse nanodomains whereas silencing marks formed large clusters (Xu et al. 2018). These observations are consistent with our previous report showing that the size of nucleosome clutches correlates with the amount of heterochromatin and nucleosome clutches become smaller and more dispersed in hyperacetylated, transcriptionally active cells (Ricci et al. 2015).

Here we carried out multi-color super-resolution imaging of histones together with DNA to define how the DNA is compacted by the nucleosome clutches and how this compaction changes after histone acetylation. Our results showed that the percentage of DNA co-localizing with nucleosomes decreased after TSA treatment. Previously, we modeled chromatin organization using a simple polymer model in which nucleosome occupancy was varied (Ricci et al. 2015). This model showed that the experimentally observed decrease in nucleosome clutch size after TSA treatment could be recapitulated through a nucleosome removal mechanism leading to lower nucleosome occupancy. Our two-color imaging data that showed a decrease in the percentage of DNA co-localizing with nucleosome clutches are consistent with the results of this synthetic model. Disruption of both histone-DNA and inter-nucleosome interactions may lead to destabilization, sliding and eviction of nucleosomes finally leading to lower DNA occupancy. Alternatively, disruption of inter-nucleosome interactions may lead to splitting apart of large nucleosome clutches, leading to opening up and release of previously not detectable nucleosome-free linker DNA. Future work is needed to discriminate between these different models: i.e. nucleosome loss from DNA versus nucleosome clutch splitting.

Further, in the super-resolution images, we could discriminate DNA associated to nucleosome clutches, which we termed “clutch” DNA. Our results suggest that the density and length-scale of clutch DNA, and hence its level of compaction, is dependent on histone tail acetylation. This result is consistent with a decrease in nucleosome clutch size after TSA treatment (Ricci et al. 2015) and shows that the level of DNA compaction is directly correlated to clutch size and nucleosome packing density of a clutch. These results are also in agreement with recent multi-color super-resolution imaging of DNA and histone modifications where H3K9ac histone modification was found to be more abundant at the less compacted regions of DNA (Xu et al. 2018), suggesting decondensation of DNA when histones are acetylated. At the molecular level, a disruption of histone-DNA interactions in hyperacetylated cells would account for the decompaction of clutch DNA observed in our experiments.

We found DNA decompaction due to acetylation to be most prominent in highly folded regions of the chromatin where clutches influence the DNA compaction of other neighboring clutches in close vicinity. It is likely that many histone tails of groups of clutches within large domains undergo hyperacetylation and hence the DNA compaction is perturbed to the highest level in these regions. Ultimately, this acetylation-dependent DNA decompaction regulates accessibility of RNA polymerase II and transcription factors to DNA regulatory sequences that may lie in these highly folded and compacted regions of the chromatin. Indeed, in fibroblasts we previously showed that RNA polymerase II is associated to the small and less dense clutches, similar to the ones generated after TSA treatment (Ricci et al. 2015). A gene likely contains multiple clutches and it would be interesting in the future to investigate if RNA polymerase II associates mostly with the decompacted DNA generated between the neighboring nucleosome clutches located in coding genes. Such visualization could ultimately be informative to study transcriptional activation. We and others (Fang et al. 2018; Kraus et al. 2017; Neguembor et al. 2018; Chen et al. 2013; Chen et al. 2018) have developed methods to image specific genomic sites, opening up the possibility to visualize DNA density and compaction at clutches located in specific regulatory regions in order to correlate function with the structure of DNA.

## Materials and Method

### Cell Culture and Sample Preparation

Human Fibroblasts (hFb) (BJ, Skin Fibroblast, American Type Culture Collection, ATCC CRL-2522) were cultured in DMEM (#41965062, Gibco) supplemented with 10% FBS (#10270106, Gibco), 1X Non-Essential Amino Acids (#11140050, Gibco), 1X Penicilin/Streptomycin (#15140122, Gibco) and 1x GlutaMax (#35050061, Gibco). For DNA labeling experiments, cells were cultured with 5μM Ethynil-deoxy-Cytdine (EdC) (#T511307, Sigma-Aldrich) for 96h previous to fixation. For hyperacetylation experiments, hFb were treated with 300nM of TrychostatinA (TSA) (#T8552 Sigma-Aldrich) in complete growth medium supplemented with 5μM EdC during the final 24h of the 96h EdC incubation before fixation. When using fiduciary markers for drift correction and 3D overlap, growth media with EdC was supplemented with 1:800 dilution of 160nm amino yellow beads (#AFP-0252-2, Spherotech) for the final 1 hour prior to fixation to permit internalization of the beads into the cells prior to fixation.

### Cell cycle analysis

hFb were grown in growth media supplemented with EdC for 96h. The cells were collected and were resuspended gently with ethanol 70% at −20C. The cells were pelleted, washed and resuspended in Propidium Iodide (Molecular Probes, #P-1304) 0.03 mg/ml, Sodium Citrate 1.1mM and RNAse A (Sigma, #R-5503) 0.3mg/mL in PBS at 4C overnight. Flow cytometry analysis was performed in a FACSCalibur (BD Biosciences). Cell cycle analysis was performed using the ModFit program.

### Staining for STORM

For imaging experiments, cells were plated on 8-well Lab-tek #1 borosilicate chambers (#155411, Nunc) at a seeding density of 20000-30000 per well for immunostaining experiments; and at a seeding density of 5000-10000 cells per well for DNA labeling experiment to allow cell proliferation and hence EdC incorporation to the genome. The cells were fixed with 4% PFA (#43368, Alfa Aesar) diluted in PBS for 15min at room temperature (RT). Then, they were permeabilized with 0.3% v/v Triton X-100 (#327371000, AcrosOrganics) in PBS for 15 min at room temperature. Cells were then blocked using 10% BSA (#9048468 Fisher Scientific) (w/v), 0.01% v/v Triton X-100 in PBS. Cells were incubated overnight with the rabbit polyclonal anti-H2B (Abcam, #1790) primary antibody diluted 1:50 in blocking buffer, at 4C with gentle rocking. Finally, cells were washed three times with blocking buffer, then incubated with 1:50 secondary antibody for STORM Imaging (see below) for 1h at room temperature. For PAINT labeling, we used the Ultivue Paint Kit (Ultivue-2). Cells were incubated for 2 h at room temperature with 1:100 of Goat-anti-Rabbit (D2) antibody diluted in Antibody Dilution Buffer. Lastly, click chemistry was performed. For click chemistry, we prepared a reaction consisting of Hepes pH8.2 150mM, Amino Guanidine 50mM (#396494, Sigma), Ascorbic Acid 100 mM (#A92902, Sigma), CuSO4 1mM, Glucose 2% (#G8270, Sigma), Glox (described in STORM imaging) 1:1000 and Alexa647 azide 10nM (#A10277, Thermo). The sample was incubated in this reaction for 30min at RT. Repeated washing was done at every step.

Secondary antibody used was donkey-anti rabbit NHS ester (Jackson ImmunoResearch) custom labeled with AF405/AF647 activator reporter dyes as previously described (Bates et al. 2007).

### STORM imaging and analysis

Imaging of H2B in single color images with spectrally different dyes was performed on a custom-built inverted microscope based on Nikon Eclipse Ti frame (Nikon Instruments). The excitation module was equipped with five excitation laser lines: 405 nm (100 mW, OBIS Coherent, CA), 488 nm (200 mW, Coherent Sapphire, CA), 561 nm (500 mW MPB Communications, Canada), 647 nm (500 mW MPB Communications, Canada) and 750nm (500 mW MPB Communications, Canada). Each laser beam power was regulated through AOMs (AA Opto Electonics MT80 A1,5 Vis) and different wavelengths were coupled into an oil immersion 1.49 NA objective (Nikon). An inclined illumination mode (Tokunaga et al. 2008) was used to obtain the images. The focus was locked through the Perfect Focus System (Nikon). The fluorescence signal was collected by the same objective and imaged onto an EMCCD camera (Andor iXon X3 DU-897, Andor Technologies). Fluorescence emitted signal was spectrally filtered by either a Quad Band beamsplitter (ZT405/488/561/647rpc-UF2, Chroma Technology) with Quad Band emission filter (ZET405/488/561/647m-TRF, Chroma), or a custom Penta Band beamsplitter (ZT405/488/561/647/752rpc-UF2) with a Penta Band Emission filter (ZET405/488/561/647-656/752m) as in (Boettiger et al. 2016). STORM and STORM+PAINT raw image data were acquired at 20 ms per frame. 488nm, 560nm, 647nm or 750nm lasers were used for exciting the reporter dye and switching it to the dark state, and a 405nm laser was used for reactivating the reporter dye. An imaging cycle was used in which one frame belonging to the activating pulse laser (405) was alternated with three frames belonging to the reporter dye. Single color imaging was performed using a previously described imaging buffer (Bates et al, 2007): 100mM Cysteamine MEA (#30070, Sigma-Aldrich), 5% Glucose (#G8270, Sigma-Aldrich, 1% Glox (0.5 mg/ml glucose oxidase, 40 mg/ml catalase (#G2133 and #C100, Sigma-Aldrich)) in PBS. A minimum of 45000 frames were obtained for every image.

For dual color, 3D SMLM images were acquired with a commercial N-STORM microscope (Nikon) equipped with a CFI HP Apochromat TIRF 100 × 1.49 oil objective, an iXon Ultra 897 camera (Andor) and a Dual View system (Photometrics DV2 housing with a T647lpxr dichroic beamsplitter from Chroma). The dual view allowed us to split the image on the full chip of the camera based on emission wavelength. 647nm laser was used to excite the DNA labelled with AlexaFluor 647 using a power density of ~3 kW/cm2. Simultaneously, in order to perform PAINT, the 560nm laser was used with ~0.8 kW/cm2 power density to excite the dye attached to the imager strand. The 405 laser was used for reactivating AlexaFluor 647 via dSTORM during acquisition. The 488 laser at ~0.1 kW/cm2 power density was used to illuminate the fiduciary beads, which were used for drift correction and chromatic alignment. The imaging cycle was composed by 19 frames of simultaneous 405 nm, 560 nm and 647 nm activation interspersed with one frame of 488 nm illumination. The yellow beads imaged with the 488nm laser were visible in both the red and orange channel, albeit dimly in the red channel. In all cases, Alexa 647 was progressively reactivated with increasing 405 nm laser power during acquisition up to a maximal power density of 0.020 kW/cm2. The imaging buffer was composed of 100mM Cysteamine MEA, 5% Glucose, 1% Glox and 0.75 nM Imager strand (I2-560 Ultivue) in Ultivue Imaging Buffer. Localizations were extracted from raw images of bead calibration, STORM and STORM+PAINT data using Insight 3 standalone software (kind gift of Bo Huang, UCSF).

The N-STORM cylindrical lens adaptor was used for STORM+PAINT data acquisition to obtain 3D localization data, as previously described (Huang et al. 2008). Briefly, calibration data was first acquired by imaging subdiffraction limit size beads (100nm Tetraspeck, ThermoScientific, T7279) in PBS adsorbed to clean glass at low enough dilution to enable single bead visualization. Using the NIS software STORM module, Z-calibration data was recorded as the microscope stage was moved in 10nm steps over a 1.6 m range and through the objective focal plane to image the elliptically shaped beads as they first elongated vertically, then horizontally. Bead localizations were extracted using the aforementioned Insight3 software through elliptical Gaussian fitting to extract horizontal and vertical width values (Wx and Wy, respectively). These widths were plotted versus the distance of the localization from the imaging focal plane, then fit with a 3rd order polynomial, as previously established (Huang et al. 2008). The coefficients of the polynomial function were input into the Insight3 software to enable assignment of an axial, *z*, position to elliptical localizations from STORM+PAINT data in addition to their lateral, *x* and *y*, positions. Rendering of DNA and histone images was performed using a summation of uniform Gaussian peaks having a fixed width of 20nm.

### STORM-PAINT Workflow

STORM localizations of DNA structure in the red 647nm channel were overlaid with PAINT localizations of histone structure in the orange 560nm channel through a defined, multistep data workflow. Firstly, the same Tetraspeck beads used for Z-calibration were utilized to define a single, 2nd order polynomial surface transfer function to overlay the two colors in *x* and *y* when imaged with the dual view. To this end, beads in 10-15 separate fields of view were imaged for 100 frames. The localizations were grouped into bead-clusters and their center identified. The bead centers from all fields of view were combined to create a pseudo-high density image map to define the registration between the orange and red channel and remove residual chromatic aberrations.

Raw STORM+PAINT image data was obtained using the aforementioned microscope hardware, including the dual view and NSTORM cylindrical lens, to record a minimum of 200,000 frames with 20ms camera exposure. Red-channel STORM localizations were extracted in 3D from the 20ms images. For the orange channel PAINT localizations, five sequential image frames were first summed together to obtain an effective 100ms camera exposure, then the 3D localizations were extracted from these summed frame images. The 5-frame summed images were used also to extract red-channel fiduciary bead localizations, and thereby improve the bead’s signal-to-background ratio, while the orange-channel fiduciary bead localizations were extracted from the raw 20ms images.

All orange-channel localizations were first overlaid upon the red channel localizations in *x* and *y* using the 2nd order polynomial surface transfer function. Next, bead localizations in each color were grouped into bead-clusters and used to extract and correct the drift trajectory for each color throughout data acquisition. Following drift correction of the datasets, the fiduciary bead positions were used to refine the lateral alignment of the two datasets in *x* and *y* using a linear affine transformation. Finally, the axial positions of the fiduciary beads were used to apply a rigid Z-translation to the orange dataset and fully overlay the 3D DNA and histone datasets. Once aligned, a 120nm slice centered near the imaging focal plane (*i.e*. Z=0) was selected for further co-structural analysis. This workflow was automated using functions developed in MATLAB version 2013a and 2016a.

### Voronoi Tesselation Analysis

Voronoi Tesselation Analysis was performed in MATLAB 2016a in a fashion similar to (Andronov et al. 2016). First, the lateral *x*,y localizations were input into the ‘delaunayTriangulation’ function, and then used to construct Voronoi polygons using the ‘Voronoidiagram’ function. Areas of the Voronoi polygons were determined from the vertices with the function ‘polyarea’. The local density in each data point was defined as the inverse value of the area of the corresponding Voronoi polygon. Voronoi polygons were visualized using the ‘patch’ function, wherein the look up table was mapped to cover 99% of the polygons; the smallest polygons being yellow (high density, greater than 0.02 nm2), larger polygons set to blue, (low density, lower than 0.001 nm2), and the largest 0.5% of polygons set to black.

### Radial density analysis

Radial density analysis was performed using a custom-written script in Python, version 2.7. The script takes as input the localizations of DNA, as directly obtained by dual-color STORM imaging, and the centroids of the H2B clusters, extracted by a previously developed Matlab script (Ricci et al. 2015). The coordinates of H2B clusters’ centroids were used as center for Voronoi tessellation. Voronoi polygons exceeding the nuclear periphery or in within the nucleoli regions were clipped to a hand-drawn mask (*voronoi_finite_polygons_2d.py function*). The DNA localizations falling within the clipped Voronoi polygon area of a particular clutch center were considered for further analysis. This procedure assigns every DNA localization to a single H2B cluster, preventing the over counting of DNA localizations during the computation. Starting from the H2B clusters’ centroid, circles of increasing radii (steps of 10 nm) were drawn and eventually bounded to the Voronoi polygon (i.e. for circles larger than the polygon, the edges of the polygon were used to clip the circle). The DNA density in within each clipped circle was calculated as the ratio between the number of DNA localizations falling within the clipped circle and the area of the disk (**Figure 2C**). To quantify the percentage of DNA localizations associated to nucleosome clutches, a clipped circle of 120 nm radius was drawn and the percentage of DNA localizations inside the disk over the total DNA localization in the polygon was calculated (**Figure 2D**). From this analysis we also quantified the percentage of H2B clusters that have more the 5 DNA localizations inside the clipped circle of 120 nm radius (**Figure 2E**).

### Statistical analysis

Graphpad Prism (v5.04) and Matlab 2016a were used for Statistical analysis. Unpaired two way Anova with Bonferroni multiple comparison test against not treated was used for Cell Cycle experiments.

DNA Voronoi density data in Figure 1D was obtained by binning the Voronoi density distributions for the six control cells and nine TSA-treated cells into 300 logarithmically spaced bins ranging from 0.1×10-9 to 0.94 nm2, which encompassed the maximal range of the 15 datasets. The median value and interquartile range for each bin was calculated and used to create figure 1D; p-value for two-sample Kolmogorov-Smirnov test between median value distributions is 0.0022.

The similarity matrix of figure 3A is calculated in MATLAB 2016a using the ‘kruskalwallis’ function to calculate p-value resulting from the Kruskal-Wallis test. This non-parametric version of ANOVA is applicable to the non-Normally distributed data we encounter and calculates the likelihood that rank means from two groups are drawn from the same distribution. To this end, the density of DNA localizations falling within pairs of search discs around H2B clusters is input into the function and the resulting p-value matrix comparing all off-diagonal search discs is shown in figure 3A. The diagonal elements were not calculated because the resulting p-value would indicate experimental reproducibility rather than provide information regarding changes in DNA density over a distance. Search discs having highly similar DNA densities give rise to larger p-values and are color-coded in yellow, whereas discs having low similarity give rise to small p-values and are color-coded in blue.

The nearest neighbor distance (NND) analysis of Figure 3B was performed by first grouping NND distances between H2B clusters within the same island (see Figure S3) into bins of 20nm spacing. For each bin, the mean cumulative DNA density (i.e. sum of DNA localizations within a search radius divided by the bounded search area) within a search radius of 70nm from the H2B clutch centroid was calculated for each dataset, four control nuclei and five TSA treated nuclei. Figure 3B was generated by plotting the average of the means across datasets within each bin ±standard deviation.

Statistical significances in panels C, D, E and H of Figure S3 were calculated using a two-sample t-test. Statistical significance in all panels of Figure S4 was calculated using a two-sample Kolmogorov-Smirnov test.

For all tests: ns *P* > 0.05, \**P* ≤ 0.05, \*\**P* ≤ 0.01, \*\*\**P* ≤ 0.001, \*\*\*\**P* ≤ 0.0001.

### Dataset selection & H2B cluster analysis

The inclined illumination utilized on our microscope systems often gave rise to illumination inhomogeneity that was visualized as long, straight areas of a nucleus having sparse localizations. Sub-regions having a more uniform illumination for both 560nm and 647nm laser lines were selected manually and used for H2B and DNA data analysis. A single sub-region was selected for each dataset.

We measured the median lateral localization precision for visualizing H2B *in situ* via DNA-PAINT (Schnitzbauer et al. 2017) methodology to be 20.6 nm (16.4, 25.0 interquartile range). The precision was determined by calculating lateral localization precision as the 2D width of a point cloud for localizations visualized in five or more sequential imaging frames (Olivier et al. 2013).

H2B cluster segmentation analysis was performed using the same algorithm previously published (Ricci et al. 2015).

STORM-DNA and PAINT-H2B images were subjected to a quality test as described next and only those images that passed the quality test were accepted for further analysis. DNA datasets were selected according to an estimate for their Nyquist sampling frequency as described previously (Legant et al. 2016). To this end, two localization densities were calculated for each dataset relative to the nuclear area measured by 1) a diffraction-limited image, and 2) binning localizations into boxes 20nm on each side. Dividing the 120nm slice thickness by this density value, and then taking the cube-root provided an estimate for the sampling frequency within a dataset. The mean ±  value across all datasets was found to be 30±2 nm/localization. Datasets utilized in DNA analysis (Figure 1C) were required be 32 nm/localization or lower for both densities (1) and (2).

For H2B, thresholds were applied following cluster analysis of an H2B-PAINT dataset using the resulting cluster metrics to determine the quality of a dataset. The three cluster metrics used to select datasets were: (i) localization occupied nuclear area (Figure S3), (ii) clusters per island (Figure S3) and (iii) the nuclear area comprising clusters. This latter metric is a ratio of the area covered by clusters compared to the area covered by localizations, where datasets having low ratios were found to be visually sparser than datasets having a higher ratio. Because the TSA treatment led to H2B reorganization, separate thresholds had to be set for control vs. treated cells. For control Cells, PAINT datasets used for analysis of H2B clusters (Figure S3 and S4) were required to have: (i) occupancy > 35%, (ii) clustered area > 15%, (iii) clusters per island > 3; for TSA-treated cells they were required to have: (i) occupancy > 25%, (ii) clustered area > 10%, (iii) clusters per island > 1.5. Dataset selection appeared robust because no datasets were found to have only one or two of the metrics above/below the thresholds. Rather, datasets removed from analysis were found to have all three metrics below the respective thresholds in all cases, while datasets included in the analysis had all three metrics well above the thresholds.

Datasets utilized in co-structure analysis of Figure 2 and Figure 3 were required to pass the selection criteria for both DNA and H2B.

## Acknowledgements

We thank Miguel Beato, Manuel Mendoza and Maria Victoria Neguembor for critical comments on the manuscript. This work was supported by European Union’s Horizon 2020 Research and Innovation Programme [CellViewer No 686637 to M.L. and M.P.C.]; the Ministerio de Economia y Competitividad y FEDER (SAF2011-28580, and BFU2015-71984 to M.P.C.), an AGAUR grant from Secretaria d’Universitats i Investigació del Departament d’Economia i Coneixement de la Generalitat de Catalunya (2014SGR1137 to M.P.C.), a “La Caixa-Severo Ochoa” pre-doctoral fellowship (to A.C.G.). The Spanish Ministry of Economy and Competitiveness, Centro de Excelencia Severo Ochoa 2013-2017 and the CERCA Programme/Generalitat de Catalunya (M.P.C).

J.J.O. acknowledges funding from the EC-Marie Sklodowska-Curie Individual Fellowship (VCSD G.A. 656873) and the EC-Marie Sklodowska-Curie COFUND action (ICFONest+ GA 609416).

## Author Contributions

J.J.O, A.C.G. conducted experiments. J.J.O and C.V. wrote software for data analysis. J.J.O, A.C.G and C.V. analyzed data. M.L. and M.P.C. conceived the idea, supervised research and wrote the manuscript. All authors read and revised the manuscript.

## References

Andronov, L., I. Orlov, Y. Lutz, J.-L. Vonesch, and B. P. Klaholz. 2016. ClusterViSu, a method for clustering of protein complexes by Voronoi tessellation in super-resolution microscopy. Scientific Reports 6:24084.

Bannister, A. J., and T. Kouzarides. 2011. Regulation of chromatin by histone modifications. Cell Research 21 (3):381–395.

Banterle, N., K. H. Bui, E. A. Lemke, and M. Beck. 2013. Fourier ring correlation as a resolution criterion for super-resolution microscopy. Journal of Structural Biology 183 (3):363–367.

Bates, W. M., B. Huang, G. T. Dempsey, and X. Zhuang. 2007. Multicolor Super-resolution Imaging with Photo-switchable Fluorescent Probes. Science 317 (5845):1749–1753.

Beliveau, B. J., A. N. Boettiger, M. S. Avendaño, R. Jungmann, R. B. McCole, E. F. Joyce, C. Kim-Kiselak, F. Bantignies, C. Y. Fonseka, J. Erceg, M. A. Hannan, H. G. Hoang, D. Colognori, J. T. Lee, W. M. Shih, P. Yin, X. Zhuang, and C.-t. Wu. 2015. Single-molecule super-resolution imaging of chromosomes and in situ haplotype visualization using Oligopaint FISH probes. Nature Communications 6:7147.

Boettiger, A. N., B. Bintu, J. R. Moffitt, S. Wang, B. J. Beliveau, G. Fudenberg, M. Imakaev, L. A. Mirny, C.-t. Wu, and X. Zhuang. 2016. Super-resolution imaging reveals distinct chromatin folding for different epigenetic states. Nature 529:418.

Chen, B., Luke A. Gilbert, Beth A. Cimini, J. Schnitzbauer, W. Zhang, G.-W. Li, J. Park, Elizabeth H. Blackburn, Jonathan S. Weissman, Lei S. Qi, and B. Huang. 2013. Dynamic Imaging of Genomic Loci in Living Human Cells by an Optimized CRISPR/Cas System. Cell 155 (7):1479–1491.

Chen, B., W. Zou, and B. Huang. 2018. CRISPR-Tag: an Efficient DNA Tagging System in Living Cells. bioRxiv.

Cho, W. K., J. H. Spille, M. Hecht, C. Lee, C. Li, V. Grube, and Cisse, II. 2018. Mediator and RNA polymerase II clusters associate in transcription-dependent condensates. Science 361 (6400):412–415.

Cisse, II, I. Izeddin, S. Z. Causse, L. Boudarene, A. Senecal, L. Muresan, C. Dugast-Darzacq, B. Hajj, M. Dahan, and X. Darzacq. 2013. Real-time dynamics of RNA polymerase II clustering in live human cells. Science 341 (6146):664–667.

Collepardo-Guevara, R., G. Portella, M. Vendruscolo, D. Frenkel, T. Schlick, and M. Orozco. 2015. Chromatin Unfolding by Epigenetic Modifications Explained by Dramatic Impairment of Internucleosome Interactions: A Multiscale Computational Study. Journal of the American Chemical Society 137 (32):10205–10215.

Cremer, T., and M. Cremer. 2010. Chromosome territories. Cold Spring Harb Perspect Biol 2 (3):a003889.

Cremer, T., M. Cremer, and C. Cremer. 2018. The 4D Nucleome: Genome Compartmentalization in an Evolutionary Context. Biochemistry (Mosc) 83 (4):313–325.

Dempsey, G. T., J. C. Vaughan, K. H. Chen, M. Bates, and X. Zhuang. 2011. Evaluation of fluorophores for optimal performance in localization-based super-resolution imaging. Nat Methods 8:1027.

Eberharter, A., and P. B. Becker. 2002. Histone acetylation: a switch between repressive and permissive chromatin: Second in review series on chromatin dynamics. EMBO Rep 3 (3):224–229.

Eltsov, M., K. M. MacLellan, K. Maeshima, A. S. Frangakis, and J. Dubochet. 2008. Analysis of cryo-electron microscopy images does not support the existence of 30-nm chromatin fibers in mitotic chromosomes in situ. Proceedings of the National Academy of Sciences 105 (50):19732–19737.

Fang, K., X. Chen, X. Li, Y. Shen, J. Sun, D. M. Czajkowsky, and Z. Shao. 2018. Super-resolution Imaging of Individual Human Subchromosomal Regions in Situ Reveals Nanoscopic Building Blocks of Higher-Order Structure. ACS Nano 12 (5):4909–4918.

Fussner, E., M. Strauss, U. Djuric, R. Li, K. Ahmed, M. Hart, J. Ellis, and D. P. Bazett-Jones. 2012. Open and closed domains in the mouse genome are configured as 10-nm chromatin fibres. EMBO Rep 13 (11):992–996.

Huang, B., W. Wang, M. Bates, and X. Zhuang. 2008. Three-Dimensional Super-Resolution Imaging by Stochastic Optical Reconstruction Microscopy. Science 319 (5864):810–813.

Jenuwein, T., and C. D. Allis. 2001. Translating the histone code. Science 293 (5532):1074–1080.

Joti, Y., T. Hikima, Y. Nishino, F. Kamada, S. Hihara, H. Takata, T. Ishikawa, and K. Maeshima. 2012. Chromosomes without a 30-nm chromatin fiber. Nucleus 3 (5):404–410.

Kraus, F., E. Miron, J. Demmerle, T. Chitiashvili, A. Budco, Q. Alle, A. Matsuda, H. Leonhardt, L. Schermelleh, and Y. Markaki. 2017. Quantitative 3D structured illumination microscopy of nuclear structures. Nature Protocols 12:1011.

Legant, W. R., L. Shao, J. B. Grimm, T. A. Brown, D. E. Milkie, B. B. Avants, L. D. Lavis, and E. Betzig. 2016. High-density three-dimensional localization microscopy across large volumes. Nat Methods 13 (4):359–365.

Levet, F., E. Hosy, A. Kechkar, C. Butler, A. Beghin, D. Choquet, and J. B. Sibarita. 2015. SR-Tesseler: a method to segment and quantify localization-based super-resolution microscopy data. Nat Methods 12 (11):1065–1071.

Li, B., M. Carey, and J. L. Workman. 2007. The Role of Chromatin during Transcription. Cell 128 (4):707–719.

Maeshima, K., S. Hihara, and M. Eltsov. 2010. Chromatin structure: does the 30-nm fibre exist in vivo? Current Opinion in Cell Biology 22 (3):291–297.

Neguembor, M. V., R. Sebastian-Perez, F. Aulicino, P. A. Gomez-Garcia, M. P. Cosma, and M. Lakadamyali. 2018. (Po)STAC (Polycistronic SunTAg modified CRISPR) enables live-cell and fixed-cell super-resolution imaging of multiple genes. Nucleic Acids Res 46 (5):e30.

Nishino, Y., M. Eltsov, Y. Joti, K. Ito, H. Takata, Y. Takahashi, S. Hihara, A. S. Frangakis, N. Imamoto, T. Ishikawa, and K. Maeshima. 2012. Human mitotic chromosomes consist predominantly of irregularly folded nucleosome fibres without a 30‐nm chromatin structure. The EMBO Journal 31 (7):1644–1653.

Olivier, N., D. Keller, V. S. Rajan, P. Gönczy, and S. Manley. 2013. Simple buffers for 3D STORM microscopy. Biomedical Optics Express 4 (6):885–899.

Ou, H. D., S. Phan, T. J. Deerinck, A. Thor, M. H. Ellisman, and C. C. O’Shea. 2017. ChromEMT: Visualizing 3D chromatin structure and compaction in interphase and mitotic cells. Science 357 (6349).

Quenet, D., J. G. McNally, and Y. Dalal. 2012. Through thick and thin: the conundrum of chromatin fibre folding in vivo. EMBO Rep 13 (11):943–944.

Raulf, A., C. K. Spahn, P. J. M. Zessin, K. Finan, S. Bernhardt, A. Heckel, and M. Heilemann. 2014. Click chemistry facilitates direct labelling and super-resolution imaging of nucleic acids and proteins. RSC Advances 4 (57):30462–30466.

Ricci, Maria A., C. Manzo, M. F. García-Parajo, M. Lakadamyali, and Maria P. Cosma. 2015. Chromatin Fibers Are Formed by Heterogeneous Groups of Nucleosomes In Vivo. Cell 160 (6):1145–1158.

Rothbart, S. B., and B. D. Strahl. 2014. Interpreting the language of histone and DNA modifications. Biochimica et Biophysica Acta (BBA) - Gene Regulatory Mechanisms 1839 (8):627–643.

Rowley, M. J., and V. G. Corces. 2018. Organizational principles of 3D genome architecture. Nat Rev Genet.

Schnitzbauer, J., M. T. Strauss, T. Schlichthaerle, F. Schueder, and R. Jungmann. 2017. Super-resolution microscopy with DNA-PAINT. Nature Protocols 12:1198.

Thoma, F., T. Koller, and A. Klug. 1979. Involvement of histone H1 in the organization of the nucleosome and of the salt-dependent superstructures of chromatin. The Journal of Cell Biology 83 (2):403–427.

Tokunaga, M., N. Imamoto, and K. Sakata-Sogawa. 2008. Highly inclined thin illumination enables clear single-molecule imaging in cells. Nat Methods 5 (2):159–161.

Toth, K. F., T. A. Knoch, M. Wachsmuth, M. Frank-Stohr, M. Stohr, C. P. Bacher, G. Muller, and K. Rippe. 2004. Trichostatin A-induced histone acetylation causes decondensation of interphase chromatin. J Cell Sci 117 (Pt 18):4277–4287.

Verdone, L., M. Caserta, and E. Di Mauro. 2005. Role of histone acetylation in the control of gene expression. Biochem Cell Biol 83 (3):344–353.

Wang, S., J. H. Su, B. J. Beliveau, B. Bintu, J. R. Moffitt, C. T. Wu, and X. Zhuang. 2016. Spatial organization of chromatin domains and compartments in single chromosomes. Science 353 (6299):598–602.

Weng, X., and J. Xiao. 2014. Spatial organization of transcription in bacterial cells. Trends Genet 30 (7):287–297.

Xu, J., H. Ma, J. Jin, S. Uttam, R. Fu, Y. Huang, and Y. Liu. 2018. Super-Resolution Imaging of Higher-Order Chromatin Structures at Different Epigenomic States in Single Mammalian Cells. Cell Rep 24 (4):873–882.

Zessin, P. J., K. Finan, and M. Heilemann. 2012. Super-resolution fluorescence imaging of chromosomal DNA. J Struct Biol 177 (2):344–348.

